# LandScape: a web application for interactive genomic summary visualization

**DOI:** 10.1101/866087

**Authors:** Wenlong Jia, Hechen Li, Shiying Li, Shuaicheng Li

**Affiliations:** Department of Computer Science, City University of Hong Kong, Kowloon Tong, Hong Kong; Department of Biology Engineering, City University of Hong Kong, Kowloon Tong, Hong Kong

## Abstract

**Summary:** Visualizing integrated-level data from genomic research remains a challenge, as it requires sufficient coding skills and experience. Here, we present LandScape^*oviz*^, a web-based application for interactive and real-time visualization of summarized genetic information. LandScape utilizes a well-designed file format that is capable of handling various data types, and offers a series of built-in functions to customize the appearance, explore results, and export high-quality diagrams that are available for publication.

**Availability and implementation:** LandScape is deployed at bio.oviz.org/demo-project/analyses/landscape for online use. Documentation and demo data are freely available on this website and GitHub (github.com/Nobel-Justin/Oviz-Bio-demo).

## 1 Introduction

The enormous development of sequencing technology has remarkably accelerated human genome research with the exponential accumulation of genomic data. Meanwhile, biological researches have analyzed trans-omics data from increasing samples. Many well-designed visualizations have been widely utilized in publications to report result findings clearly and thoroughly^1,2^. Recent cancer genome studies generally analyze genomic data from hundreds of tumors and compare them among a series of attributes, such as mutation burden, mutated genes, and biological pathways, which is commonly demonstrated via the well-known ‘Landscape’ figures. The Landscape figure and its variants have been prevalently applied in studies with a large cohort of samples as it is perfect for summarizing the interested multi-layer property patterns of all samples at once^3,4^. The implementation of these figures remains a challenge to biology researchers as it requires sufficient programming skills and needs much effort to fine-tune for publication. Additionally, figures generated by existing software packages are static, which makes interactive data exploration impossible.

Here, we present LandScape^*oviz*^, a web-based application for interactive and real-time visualization of integrative multi-layer data of batch samples. LandScape utilizes a well-designed format to handle multiple data types, and offers a series of built-in functions to customize the display, explore results, and generate high-quality diagrams.

## 2 Implementation

LandScape is deployed on the Oviz-Bio online platform, which is constructed with a back-end written in Ruby on Rails and an in-house visualization language (Oviz) compatible with both HTML5 Canvas and Scalable Vector Graphics (SVG). Oviz has been utilized in developing a series of common visualizations in data science (chart.oviz.org). By design, LandScape comprises some constant panels (histograms and the gene panel) and additional user-defined panels to present more information, such as age, gender, and clinical histology. Visualization is automatically generated according to the file input in the comma-separated values (csv) format.

## 3 Input file format

We designed a user-friendly and flexible format for the csv input file. The header of the csv file includes reserved keys (e.g., ‘SampleID’), keys with reserved prefixes (e.g., these for gene panel and histogram attributes), and other user-defined keys. Users could add new custom column names in the header line to supply the data for additional panels. Basically, the content of the csv file could be considered as a matrix with rows and columns representing samples and attributes, respectively. We also prepared several demo csv files to reproduce the LandScape figures used in several highly cited cancer studies^3–8^.

## 4 Constant panels of LandScape

The constant part consists of histograms and the gene panel (Fig. 1A-F). LandScape allows multiple stacked histograms to be shown at the top of the visualization (Fig. 1A-B), illustrating the distribution of sample attributes. Each attribute displayed in histograms is defined as a column name in the header of the csv input file, for instance, a header key ‘ht1 Missense’ indicates that certain property (e.g., mutation density) of the missense type mutations is an attribute displayed in the first histogram. The gene panel displays the alteration status in a gene list using a matrix style (Fig. 1C), which is usually used to illustrate significantly mutated genes in cancers. Gene names are stated in the header with reserved prefix (‘g_‘), e.g., ‘g_TP53’ for cancer gene *TP53*. Sample frequency of mutation types in each gene is calculated and displayed by the horizontal histogram at the left side of gene-panel (Fig. 1D). For scenarios such as harbouring double mutations, LandScape allows showing multiple alteration types of one given gene in a single sample (Fig. 1C). Generally, in cancer research, genes are always annotated with more information, such as KEGG pathway, GO ontology, additional comments, and *P*-value or *Q*-value representing their mutated significance in batch samples. These information (if given) are shown at the right side of gene-panel as tags, matrices or histograms (Fig. 1E-F).

**Figure 1.**
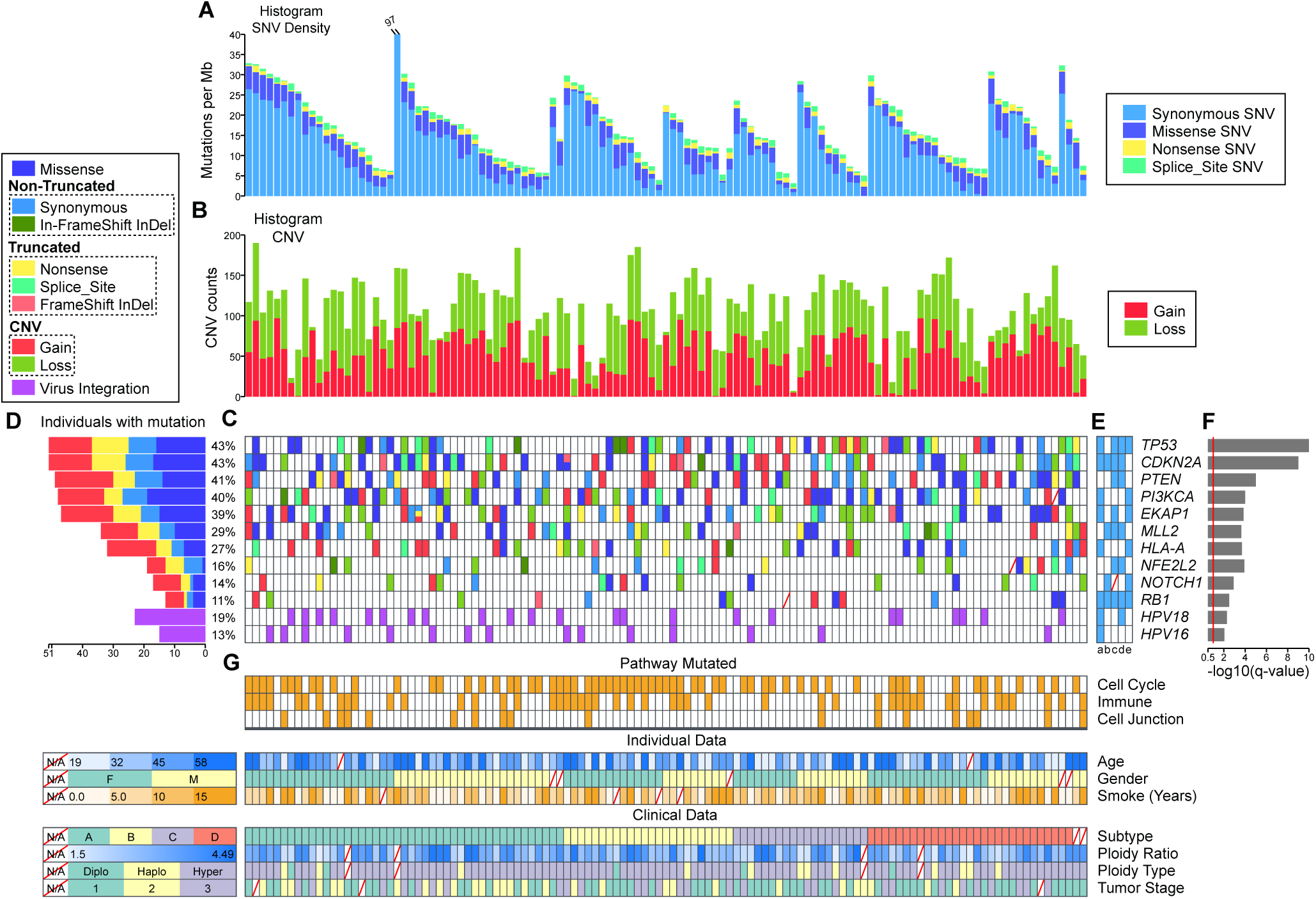
Demo representation of LandScape visualization. All panels are aligned with vertical tracks representing 119 samples, which are sorted in order by ‘Subtype’ (Clinical Data), ‘Gender’ (Individual Data), and ‘Mutation density’ (top histogram). (**A**) Histogram of somatic SNV mutation density. (**B**) Additional histogram of CNV counts. (**C**) Color-coded mutation types of SMGs. (**D**) Overall mutation frequency of SMGs. Note that certain mutation types are grouped into category (dashed frame in legend) whose color follows the first included mutation type. (**E**) Comments on genes. The comment values could be Boolean or numeric. (**F**) *Q*-value of SMGs. This panel could also display *P*-value, KEGG pathway and GO ontology of SMGs via sidebar options. (**G**) Additional panels for mutated pathway status, individual and clinical data of samples. SNV, single nucleotide variant; InDel, insertion and deletion; CNV, copy number variant; SMGs, significantly mutated genes.

## 5 Additional panels of LandScape

The additional panels is designed to show information on multiple aspects (Fig. 1G), such as patient clinical data, and mutated status at the biological pathway level. Users could add columns in csv input with self-defined header keys following the format ‘*PanelName_AttributeName*’, e.g., ‘MetaData_Age’ for patient age, and ‘Pathway_CellCycle’ for the cell cycle pathway. LandScape summarizes value types in each column and automatically applies appropriate display styles for string/boolean (classification) and numeric values (gradient color), respectively. Furthermore, users could also choose the desired value type manually and adjust details of the groups, such as value ranges for numeric values in the sidebar (e.g., ‘Age’ in Fig. 1G). LandScape displays additional panels in the order as they appear in the input.

## 6 Interactions and customization

LandScape provides many interactive built-in functions for users to explore the data in multiple aspects. Basically, when the mouse cursor moves, columns and rows will highlight to assist the aligning especially for the large sample cohort, and corresponding tooltips will appear to show necessary information of object. Genes and pathways are linked to the genecards^9^, KEGG pathway^10^, and GO ontology databases^11^, respectively. Real-time visualization can be saved in SVG file with either dark or light theme. The sidebar offers copious options to adjust the figure structure and fine-tune the appearance. Options are categorized into sections according to their functions. Users could reorder the samples with the combination of multiple criteria, including sample name, histogram values, and attributes in additional panels. LandScape also provides the well-known bisection method, which sorts samples hierarchically according to the mutually exclusive mutated genes^3, 5, 12^. Moreover, the mutation types could be grouped, for instance, the synonymous mutation is always considered as non-mutated as it causes no protein changes^3^. Besides, user could manually reorder the histogram stacks and rows in each panel, rename histograms and panels, change maximum value and labels of axis, customize the color for each object in each panel, select interested information to display at the right side of gene panel, and choose display styles for attributes with numeric values.

## 7 Conclusion

LandScape provides real-time visualization with a well-designed graphical interface and functions for the exploration of summarized data in genetic studies. The input data format is not only simple to learn but also flexible for future extension. With its versatile displaying options, LandScape will help users generate customized publication figures with minimum coding efforts. LandScape successfully reproduced figures from a series of highly cited publications, which proves its availability and practicability. More functions will be implemented as the visualizer kernel (Oviz) upgrades.

## Funding

This work was supported by the GRF Research Projects 9042348 (CityU 11257316).

## Conflict of Interest

none declared.

## Notes

#### Summary of Updates

Abstract revised; Funding statement added; Conflict of Interest declared.

https://bio.oviz.org/demo-project/analyses/landscape

https://github.com/Nobel-Justin/Oviz-Bio-demo

